# The TMRCA of general genealogies in populations with deterministically varying size

**DOI:** 10.1101/2024.09.19.613917

**Authors:** Alejandro H. Wences, Lizbeth Peñaloza, Matthias Steinrücken, Arno Siri-Jégousse

## Abstract

We study the time to the most recent common ancestor (TMRCA) of a sample of finite size in a wide class of genealogical models for populations with deterministically varying size. This is made possible by recently developed results on inhomogeneous phase-type random variables, allowing us to obtain the density and the moments of the TMRCA of time-dependent coalescent processes in terms of matrix formulas. We also provide matrix simplifications permitting a more straightforward calculation. With these results, the TMRCA provides an explanatory variable to distinguish different evolutionary scenarios, and to infer model parameters.

## 1 Introduction

The general theory of coalescent processes aims to provide a rigorous mathematical framework that can be used to model natural phenomena where a collection of particles fuses together as the system evolves over time. It has a variety of applications in different disciplines, such as physics and biology. In biology, particularly in the field of population genetics, it is used to model the parental relationships of a given sample or population, as we trace the ancestry of individuals backward in time, thus constructing a genealogical tree. In this setting, the coalescence of particles occurs at the time when a group of individuals has a common ancestor in the past. Once we have a suitable coalescent model for the genealogy of a population, we can employ mathematical tools to tackle biological questions, such as determining the time needed to reach the most recent common ancestor of the sample or population (TMRCA), the expected genetic diversity for neutral positions of the genome, or whether natural selection has played an important role in the evolution of the population [18, 25, 46].

In mathematical population genetics, the study of coalescent processes focuses on two main questions. First, a lot of effort is dedicated to establishing equivalence between coalescent processes and corresponding population models with various biological assumptions, such as constant or varying population size, the presence of mutations, the strength of natural selection, the effect of genetic drift, spatial constraints, as well as dormancy/latency mechanisms as in virus populations. The motivation behind this effort is that once a coalescent model is inferred for the genealogy of a population from genetic data, this equivalence allows for the inference of the evolutionary forces that significantly influenced the dynamics of the population. One of the first coalescent models, Kingman’s coalescent, was established as the null model for neutral genealogies of populations where the variance of the offspring distribution is much smaller than the population size, evolving at equilibrium, that is, the population size stays constant [32, 39]. In more recent years, the Bolthausen-Sznitman coalescent has emerged as an alternative model for the genealogy of constant-size populations that nonetheless are subject to the effect of natural selection [11, 40, 12, 41]. Here, a burst of reproductive success of individuals carrying a beneficial genotype leads to multiple mergers in the corresponding genealogy. The genealogies of populations with stochastically varying population size, or evolving in a random environment, have also been addressed; for example, neutral populations undergoing recurrent (i.e. i.i.d. across generations) bottlenecks were studied in [29, 20, 19, 43] for both high and low-variance reproductive laws. Also, neutral populations evolving under deterministically-varying population size and with low-variance reproductive laws were studied in the seminal work of [23]; the coalescent that describes the genealogies of these populations is a time-inhomogeneous coalescent process that can be expressed as a deterministically time-changed Kingman’s coalescent.

The second main question addressed by the study of coalescent processes concerns the theoretical characterization of different functionals of coalescent processes such as the tree height, the total tree length, or the size of external and internal branches. This provides inference tools for distinct aspects of the evolutionary past of a population, such as the forces at play throughout its history, the presence of bottlenecks, the TMRCA, etc. In this work, we are interested in the study of the density and moments of the TMRCA for time-inhomogeneous coalescent processes describing the genealogies of populations evolving under deterministically-varying population size. This functional, apart from being interesting as a mathematical object in its own right, is very useful as a step-variable in applications such as the inference of demographic history (see, e.g. [28, 47]) or in the computation of the expected SFS [45]. This variable was previously analyzed for particular examples such as Kingman’s coalescence with general time-change in [23] (first moment), but also in [50] (second moment), and in [14] (any moment). For general coalescent models with piecewise constant time-change, the first moment was established in [45]. All these methods are hard to generalize due to analytical difficulties caused by the time dependence and combinatorial issues when trying to consider more general models, higher moments, or their density function. Here, we provide a new technique based on inhomogeneous phase-type theory, developed in [2], to efficiently obtain any moment and the density of the TMRCA for general Markovian genealogical models with a deterministic time-change function.

The phase-type theory, in its inhomogeneous setting, is less flexible than its homogeneous version developed in [27]. Hence it does not provide tractable formulas for other statistics such as total length or site frequency spectrum. Our techniques give considerable analytical advantage to the use of the TMRCA in applications in population genetics, specially when the population under consideration has a varying size. Indeed, we show how explicit analytical formulas for the likelihood function of an i.i.d. sample of TMRCAs can be effectively used to infer parameters of evolutionary models: We demonstrate its use for the inference of the rate of growth in exponentially growing populations, as well as the characterization of the shape of the underlying genealogy, through parameter estimation via maximum likelihood estimators (MLE).

## 2 The model

*Time-changed coalescents*. We consider the general class of deterministically time-changed Ξ-coalescents characterized by a finite measure Ξ on the simplex 𝒫 _[0,1]_ := {(*p*_*i*_)_*i≥*1_ : *∀p*_*i*_ *≥* 0 for all *i ≥* 1, and 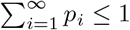,} and a deterministic time-change function *g*^−1^(*t*) —the notation was chosen to coincide with that in [2]— of the form

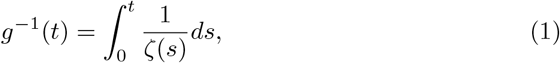

with *ζ*(*t*) *>* 0 for all *t* > 0. These processes are of the form (Π(*g*^−1^(*t*)))_*t≥*0_, where (Π(*t*))_*t≥*0_ is a time-homogeneous Ξ-coalescent process. The time-change has the effect of scaling the coalescent rates by the function of time 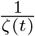. They arise as the genealogy of a sample of individuals from a population whose total size evolves deterministically over time. Namely, going backward from the time when the sample of individuals is taken —which corresponds to time 0 in the coalescent—, the dynamics of the total size of the population is described by the function *ζ*(*t*) (see e.g. [23]).

We now describe this class of processes in more detail: For a given sample size (or number of particles) *n* > 0, these processes are pure-jump time-inhomogeneous

Markov processes with state space the set of partitions of [*n*] := {1, …, *n*}. The states represent configurations of blocks, and the initial state is the partition into singletons {{1}, …, {*n*}}. Then, as the coalescent progresses —going backward in time in the population—, blocks representing ancestral lineages find common ancestors, leading to coagulation events that form new bigger blocks and thus coarser partitions of [*n*]. The absorbing state {1, *· · ·, n*} is eventually reached, where there is only one lineage that is ancestral to all the initial particles. The time it takes to reach the absorbing state corresponds to the TMRCA of the *n* sampled lineages.

Specifically, at time *t ≥* 0 a jump directed by ***p*** = (*p*_*i*_)_*i≥*1_ occurs at rate 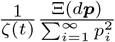. At such an event, each active lineage (i.e. block of the current partition) is independently and uniformly placed in the interval [0, 1], which is divided into subintervals of lengths (*p*_*i*_)_*i≥*1_. The new state of the system (a coarser partition of [*n*]) is constructed by merging (coalescing) all the lineages that fall into the same subinterval. Note that, when 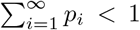, the lineages falling outside of any subinterval do not participate in any coagulation (this is Kingman’s paintbox construction, see e.g. [4]). In general, Ξ-coalescents are called simultaneous multiple merger coalescents, since more than one group of lineages can coalesce into distinct ancestral lineages at the same time as soon as at least two values in (*p*_*i*_)_*i≥*1_ are non-zero.

When the measure Ξ is supported on the set {(*p*_*i*_)_*i≥*1_ : *p*_1_ *∈* (0, 1], and *p*_*i*_ = 0 *∀i ≥* 2} the corresponding coalescent is called a —time-inhomogeneous— multiple merger coalescent, since only one group of lineages coalesces at a time. In this case, Λ denotes the push-forward measure of Ξ on [0, 1] after projecting into the *p*_1_ coordinate. Similar to the Ξ-coalescent, any lineages that are placed in the unit interval above *p*_1_ for a specific coalescent event do not participate in the event. Such lineages are also referred to as “dust.” Finally, our results also apply to coalescent processes with Kingman’s dynamics, in which every pair of blocks independently coalesces at an additional rate of 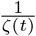.

Established coalescence processes/measures in the literature and their associated population dynamics are:

- Kingman’s coalescent (Ξ = 0): For most populations evolving at equilibrium that have an offspring distribution with variance much smaller than the population size. Here, only one pair of ancestral lineages can coalesce at a given time.
- Beta coalescents (Λ *∼* `Beta`(2 − *α, α*), 1 ≤ *α* < 2): For populations with skewed offspring distribution [42] or selection [11, 40, 12, 41]; this class includes as limit cases the Kingman coalescent (*α →* 2, neutral evolution) and the Bolthausen-Sznitman coalescent (*α* = 1, strong selection).
- Psi coalescent (Λ*∼ δ*_*ψ*_, 0 < *ψ* ≤ 1): For populations with skewed offspring distribution [13]. Here, at a given coalescent event, each active lineage participates in the coalescent event with probability *ψ*.
- Symmetric coalescent (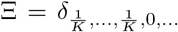 with *K* random, see Example 2 in Section 5 for more details): For populations with recurrent drastic bottlenecks [20, 19].
- Ξ^*β*^-coalescent: The Ξ^*β*^-coalescent approximates the ancestry of Cannings models with seed bank effects, where 0 < *β* < ∞ is the expected number of generations that separates an individual from its ancestor. The paintbox construction of the Ξ^*β*^-coalescent is obtained by dividing all subintervals in the paintbox construction of the associated Ξ-coalescent by *β* [22].
- Beta-Ξ-coalescent (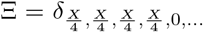 with *X ∼* `Beta`(2 − *α, α*)): Modeling diploid reproduction populations with large family sizes [9, 6]. In diploid models, there can be up to four simultaneous coalescent events per large family event.

In population genetics, the time-change function *ζ*(*t*) captures the dynamics of the effective population size, resulting in a scaling of the coalescent rate by 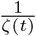: The coalescent is accelerated in small populations, whereas it slows down in larger populations. This scaling was originally introduced in the works of Griffiths and Tavaré [23] (theoretical biology), and Möhle [38], Kaj and Krone [29] (probability). For more details, see also [17]. Common examples of time-changes functions *ζ*(*t*) are

- Exponential growth: *ζ*(*t*) = *e*^−*ρt*^, with *ρ* > 0. The population grows exponentially forward in time at a rate of *ρ* [24].
- Frequent bottlenecks: *ζ*(*t*) = 1 + *ε* sin(*ωt*). The population size fluctuates around 1, with amplitude *ϵ* and frequency *ω* [14].
- Piecewise constant function 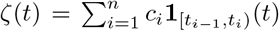, with *c*_*i*_ > 0 *∀i*, and *t*_0_ = 0 < *t*_1_ < … < *t*_*n*_ = *∞*. The population is of constant size *c*_*i*_ during the interval [*t*_*i*−1_, *t*_*i*_) [35, 45].
- Piecewise exponential function 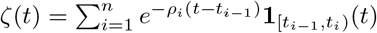, with *ρ*_*i*_ *∈*ℝ *∀i*, and *t*_0_ = 0 < *t*_1_ *<* … *< t*_*n*_ = ∞. Exponential growth or decline in distinct intervals. The rates *ρ*_*i*_ are often matched such that the sizes agree at the change-points *t*_*i*_ [5].

We can also consider variations of coalescent processes representing genealogies of populations with more complex evolutionary scenarios, such as certain types of structure [34] or seed bank coalescent [21]. Our method can be applied in any scenario where the process describing the number of lineages is a time-rescaled homogeneous Markov chain with a finite state space. We note that this does not readily include genealogical processes with branching, such as the ancestral selection graph [33], for example. This is due to the fact that the maximal number of lineages is not known a-priori, and thus the respective state space is not finite. In special cases, a version restricted to finite states might still provide a good approximation.

## 3 Methods

Our approach will make use of phase-type distributions, which define a class of random variables including sums and mixtures of exponentials. They are commonly defined as the time to absorption of a continuous-time Markov chain. Methods based on phase-type distributions have been used in biology and medicine [1, 36, 15] and, more recently, in population genetics [27, 26], in particular, to examine balancing selection [49], or seed bank dynamics [8, 21].

As shown in [27] for time-homogeneous Markovian genealogies, some of the most important statistics on coalescent processes such as the TMRCA and the total branch length can be cast into the phase-type framework, leading to explicit expressions for their density and their moments in terms of a suitable chosen rate matrix of a Markov process with absorption. In Section 4, building on previous work [2], we provide simple formulas for the density and the moments of the TMRCA in the time-inhomogeneous setting for a variety of models.

*Preliminaries on phase-type distributions*. Consider a general Markov chain with *n* states, say {1, …, *n*}, including an absorbing state, and time-inhomogeneous transition matrix of size *n × n* given by

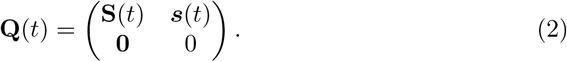

Here we follow the usual convention that the initial state has index 1, and the absorbing state has index *n*. Thus, the last row of zeros in this matrix corresponds to the jump rates of the absorbing state; whereas the column vector ***s***(*t*) of size *n* − 1 gives the infinitesimal jump rates at time *t* from any non-absorbing state to the absorbing state. Moreover, the matrix **S**(*t*) of size (*n*− 1) *×* (*n*− 1) encodes the jump rates, at time *t*, between non-absorbing states; i.e. **S**_*i,j*_(*t*) is the jump rate from a non-absorbing state with index *i ∈* {1, …, *n* − 1} to another non-absorbing state with index *j ∈* {1, …, *n* − 1}. Finally, 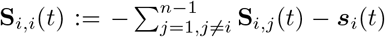. The whole matrix **Q**(*t*) can thus be recovered from **S**(*t*), and **Q**(*t*) is a valid Markov transition rate matrix. This can then be used to define [2, Definition 2.1]:

**Definition 3.1** (Inhomogeneous phase-type distribution). *The random time τ when the Markov process with rate matrix* **Q**(*t*) *reaches its absorbing state is called an* ***inhomogeneous phase-type distributed*** *random variable with parameter* **S**(*t*).

In our setting the absorbing Markov chain will typically be the block-counting process associated with a time-inhomogeneous coalescent process. This process is a death process that starts with *n* initial lineages or blocks (corresponding to state 1 in **S**(*t*)). Under the dynamics of the coalescent, lineages are successively merged, thus decreasing the number of blocks, until the process reaches the absorbing state where there is only one lineage left that is ancestral to all particles (corresponding to index *n* in **S**(*t*)): the MRCA of the underlying population. This setting, however, can be easily generalized to more complex dynamics for the number of blocks, for instance by distinguishing between “dormant” or inactive and active blocks. Indeed, we need only ensure that the first index in the matrix **Q**(*t*) is the initial state, and the last index is the absorbing state of the system. We will present such a generalization in Example 3 in Section 5 where we consider time-inhomogeneous coalescent processes with dormancy.

Our main result is that the density, the Laplace transform and the moments of *τ* can be computed in terms of the matrix **S**(*t*) and the vector ***s***(*t*).

## 4 Results

In this section we derive results for the TMRCA of general Ξ-coalescent models with time-change function *g*^−1^(*t*), introduced in Section 2. As explained above, instead of considering the whole partitioned-valued process, we need only consider the block-counting process of the time-homogeneous coalescent model. The rate matrix of the block-counting process for the time-changed coalescent is given by

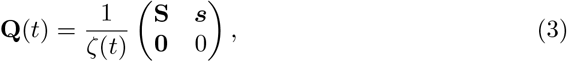

where *ζ*(*t*) is a function satisfying *ζ*(*t*) > 0 for all *t* > 0, **S**_*i,j*_ is the rate of the jump decreasing the number of lineages from *n* − *i* + 1 to *n* − *j* + 1, and ***s***_*i*_ is the rate of jumping from *n* − *i* + 1 lineages to the absorbing state with one remaining lineage.

Recall Kingman’s paintbox construction described in section 2 for the dynamics of the Ξ-coalescent but this time without time-change (e.g. *ζ*(*t*) *≡* 1). Assume that at time *t* there are *n* − *i* + 1 blocks. To compute the rate **S**_*ij*_ at which the number of blocks jumps from *n* − *i* + 1 to *n* − *j* + 1, consider the event that *k ≥* 1 groups of blocks of respective sizes *b*_1_ *≥* 2, *…, b*_*k*_ *≥* 2 coalesce simultaneously, with *n* − *i*+1− (*b*_1_+*…* +*b*_*k*_)+*k* = *n*− *j* +1. With the notation 1 = *b*_*k*+1_ = *…*= *b*_*n*−*j*+1−*k*_ we have, summing over all such possibilities:

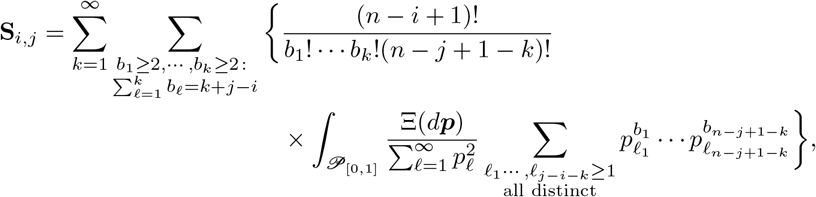

where empty sums are set to 0. Note that the factor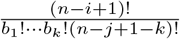 accounts for choosing a combination of *k ≥* 1 groups of blocks of sizes *b*_1_, *…, b*_*k*_. The sum inside the integral computes the probability, in the paint-box construction, that the chosen groups of blocks coalesce simultaneously by having their respective uniform random variables fall on the same interval of respective sizes 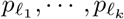; and also that the rest of the blocks do not participate in any coagulation event, by having their respective uniform random variables fall on distinct intervals of sizes 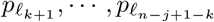. This is computed for all possible choices of 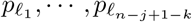 in ***p***; where we recall that ***p*** arrives with intensity 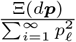.

The above expression simplifies for Λ-coalescents. Taking *b* such that *n* − *i* + 1 − *b* + 1 = *n* − *j* + 1, we obtain

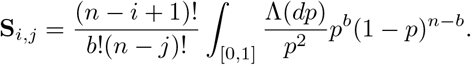

Note that since the number of lineages can only decrease, the matrix **S** and thus **Q**(*t*) is upper triangular. From Definition 3.1, it follows that the random time *T*_MRCA_ when the most recent common ancestor —the absorbing state with index *n* in **S**— is reached has an inhomogeneous phase-type distribution.

This special case is simpler than the general case and is treated in Theorems 2.8 and 2.9 of [2], which we restate in our notation as:

### Theorem 4.1.

*The random time T*_*MRCA*_ *has the same distribution as g*(*X*) *where X ∼ PH*(**S**) *(homogeneous) and the inverse of g is given by* (1) *above. In particular, its density function is given by*

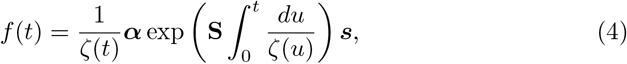

*where* ***α*** = (1, 0, …). *Also, the k-th moment of T*_*MRCA*_ *is given by*

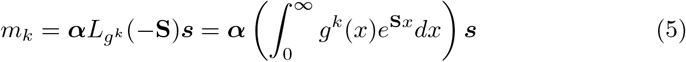

*where* 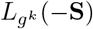 *denotes the matrix-valued Laplace transform of g*^*k*^, *parametrized by the matrix* −**S**.

The *matrix-valued Laplace transform* in equation (5) is difficult to implement. In Theorem 4.2, we provide a modification of (5) involving the matrix-valued Laplace transform applied to each eigenvalue, which significantly eases the computation. This works when the genealogical model considered is given by any time-inhomogeneous Ξ-coalescent (see also Remark 1).

### Theorem 4.2.

*For any* Ξ*-coalescent starting with n particles there exists a matrix* **P** *≡* **P**_Ξ,*n*_ *and a vector* ***s*** *≡* ***s***_Ξ,*n*_ *such that for any deterministic time-change function ζ*(*t*) *and g*(*t*) *as in* (1) *we have, for the density f of T*_*MRCA*_,

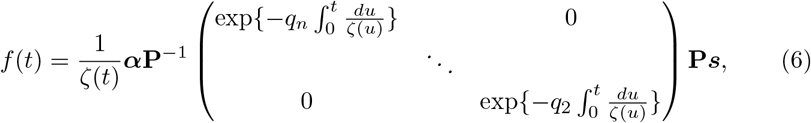

*where q*_*j*_ *is the total jump rate of the* Ξ*-coalescent when there are j particles. Similarly, for the k-th moment of T*_*MRCA*_,

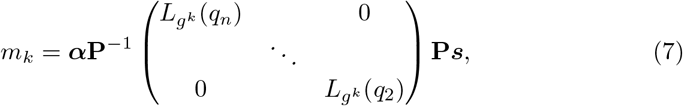

where 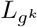 denotes the regular scalar Laplace transform.

*Proof*. Consider the Markov chain rate matrix **Q** where **Q**_*i,j*_ is the rate at which the (homogeneous) block-counting process jumps from *n*− *i* + 1 to *n* − *j* + 1, with 1 *≤ i < j ≤ n*. Note that the corresponding matrix **S** defined in equation (3) is upper triangular and diagonalizable. In particular,

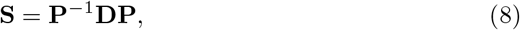

where the rows of **P** are the (left) eigenvectors of **S** and **D**_*i,i*_ = **S**_*i,i*_ for 1 *≤ i < n*, since the eigenvalues of **S** are {**S**_1,1_, …, **S**_*n*−1,*n*−1_}. Then, since ***α*** = (1, 0, …, 0),

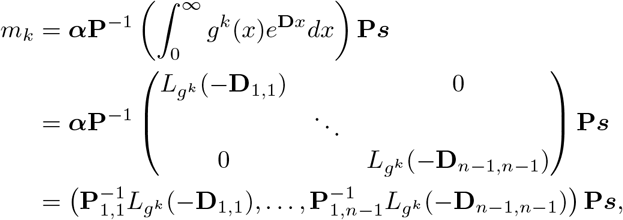

and, similarly,

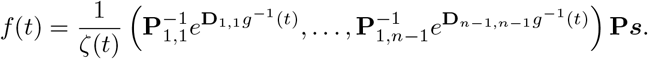

The statement of the theorem follows from the observation that **D**_*i,i*_ = **S**_*i,i*_ = −*q*_*n*−*i*+1_.

### Remark 1.

*The above proof rests solely on the (left) eigenvalue decomposition of* **S** *as in* (8). *Thus, equations* (6) *and* (7) *remain valid for any inhomogeneous phase-type distribution with transition matrix of the form* (3) *satisfying* (8), *as long as the corresponding eigenvectors* **P** *and eigenvalues (which will replace* (−*q*_1_, *…*, − *q*_*n*_) *in (6) and* (7)*) are used*.

The following lemma aims at easing the computation of the vector 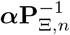.

### Lemma 4.3.

*Let* **P**_Ξ,*n*_ *be as in Theorem 4*.*2*.

1. *The matrix* **P**_Ξ,*n*_ *can be obtained by removing the first row and the first column of* **P**_Ξ,*n*+1_.
2. *The* (1, *i*)*-th entry of* 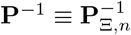 *is given by*

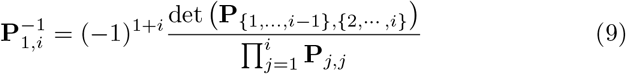

*where, for I, J ⊂* [*n* − 1], *we have used the notation* **P**_*I,J*_ *to denote the matrix* (**P**_*i,j*_)_*i∈I,j∈J*_.

*Proof*. Item 1) follows from the fact that **S** *≡* **S**_*n*_ is upper triangular for every *n* and that **S**_*n*_ can be obtained from **S**_*n*+1_ by removing its first row and column.

To prove item 2), note that if 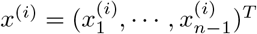 is the solution to

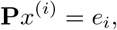

where *e*_*i*_ is the *i*-th unit vector, then, by Cramer’s rule,

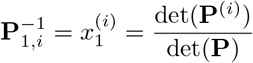

where **P**^(*i*)^ is constructed from **P** by replacing the first column by the vector *e*_*i*_. Computing the determinant of **P**^(*i*)^ along the first column using the Laplace expansion then gives

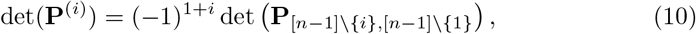

since all elements of the first column are zero, except the *i*-th entry, which is equal to one.

Since **S** is upper-triangular, the matrix **P** of its left eigenvectors is upper-triangular, and so is **P**_{*i*+1,*···, n*−1},{*i*+1,*···, n*−1}_. This block structure for the determinant on the right-hand side of (10) implies

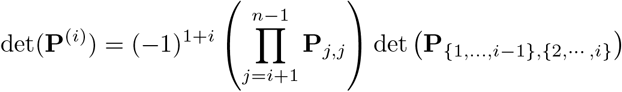

And

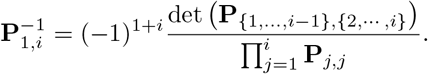

Equation (9) can be evaluated for all 1 *≤ i ≤ n* − 1 efficiently using a dynamic program that reuses computations for *i* − 1 in the computations for *i*. To this end, define

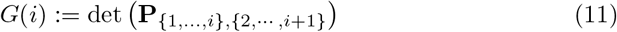

and

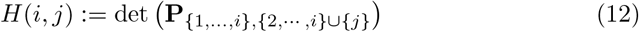

for 1 *≤ i < n* − 1 and *i* + 1 *< j ≤ n* − 1. Now note that for all *i* and *j*, the matrices in the definitions (11) and (12) only have two non-zero entries in the last row, specifically, in the last two columns. We can thus compute the determinants along the last row using the Laplace expansion to show that the recurrence relations

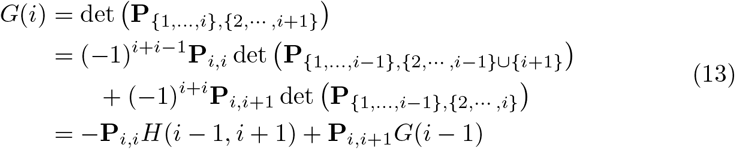

and

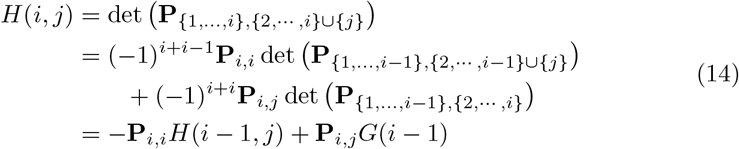

hold. The functions *G*(*i*) and *H*(*i, j*) for all *j* can then be computed iteratively, starting with *i* = 1 and incrementing *i* by 1 until *i* = *n* − 2, reusing the values of *G*(*·*) and *H*(*·,·*) from the previous step. Once *G*(*i*) is computed for all *i*, equation (9) can be readily evaluated for all *i*. Note that we did observe improved numerically stability when absorbing all but the highest factor of the product in the denominator of equation (9) into **P** in the numerator by dividing each row *i* with the respective value **P**_*i,i*_ on the diagonal.

## 5 Applications

The purpose of this section is two-fold: 1) to show that the TMRCA can be used to distinguish between different evolutionary models for a population; 2) to demonstrate the use of our analytic formulas for inference problems in population genetics. In practice, the TMRCA can be estimated from genetic data, for example, by using software to infer genealogies [44, 31]. Of particular note here is `tsinfer+tsdate` [48], which can perform inference of the TMRCA without a priori assumptions about the population size function *ζ*(*t*).

In the following Example 1 we use explicit expressions for the likelihood function, and its gradient, of a sample of *N* independent measurements of the TMRCA of *n* individuals in order to obtain maximum likelihood estimates (MLE) of population parameters. In general, one can consider growth functions 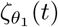 parametrized by *θ*_1_, and coagulation matrices 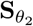 parametrized by *θ*_2_. The likelihood function, corresponding to a sample 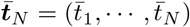 of TMRCA’s can then be written as

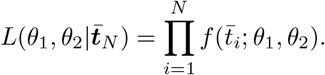

The latter expression can be used in gradient-based optimization algorithms to obtain the MLE. The derivatives of the density function *f* (*t*; *θ*_1_, *θ*_2_) with respect to these parameters, from which the gradient of 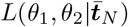 is easily obtained, are:

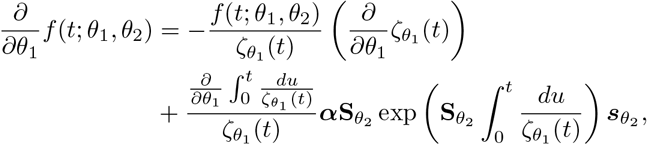

And

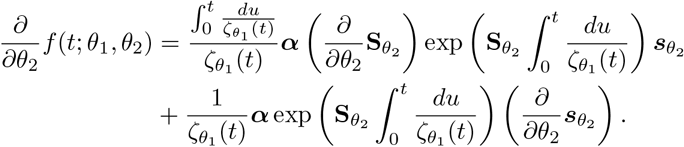

Here we recall that, for each *θ*_2_, the computation of the exponential 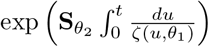 can be decomposed as in Theorem 4.2. In the following examples, we revisit some of the models introduced in Section 2, to demonstrate how the TMRCA can be used to distinguish between different models and to infer model parameters.

*Example 1: Multiple merger coalescents with exponential growth*. For the special case of exponential growth with rate *ρ* forward-in-time, the time-scale function is given by *ζ*(*t*) = *e*^−*ρt*^, and we obtain *g*^−1^(*x*) = *ρ*^−1^(*e*^*ρx*^ − 1) and *g*(*x*) = *ρ*^−1^ log(1 + *ρx*). Thus

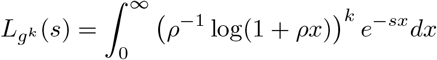

and, for any Ξ-coalescent,

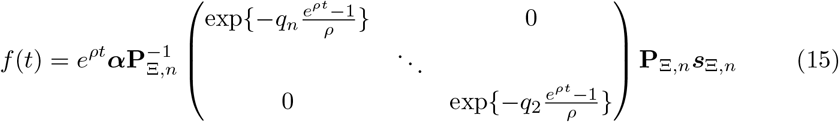

follows.

We now focus on the subfamily of the Beta coalescents, with exponential growth (see Section 2). In Figure 1, we show the density of *T*_MRCA_ for Kingman’s coalescent and the Bolthausen-Sznitman coalescent (*α* = 1) with *n* = 30 for different values of *ρ*. In the same scenarios, Figure 2 depicts the respective moments *m*_*k*_ for different values of *k*.

**Figure 1.**
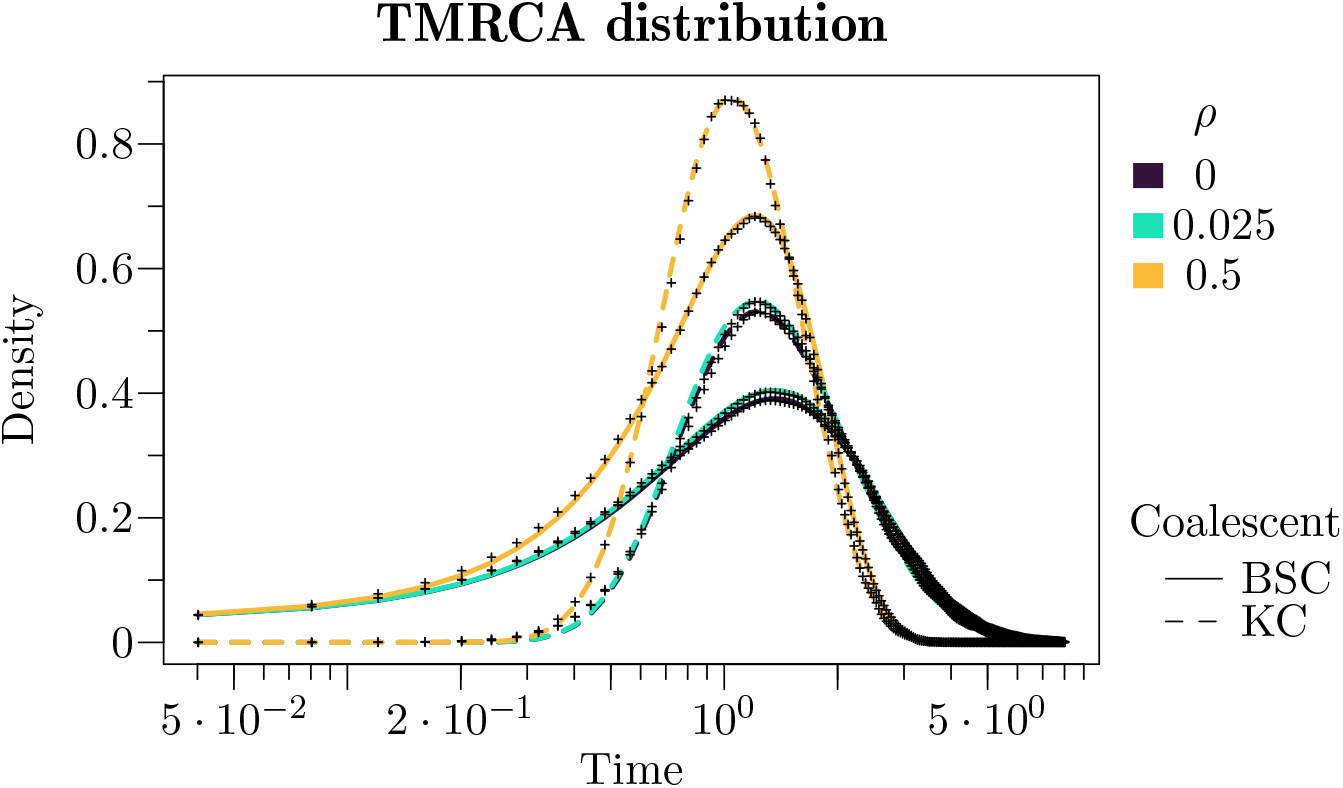
The density of *T*_MRCA_ for different choices of *ρ* in the exponential growth model, for Kingman’s (KC) and the Bolthausen-Sznitman coalescent (BSC) with *n* = 30. We also provide a visual verification comparing the values obtained from our analytical formulas against densities estimated from 40,000 simulated replicates (indicated by plus signs).

**Figure 2.**
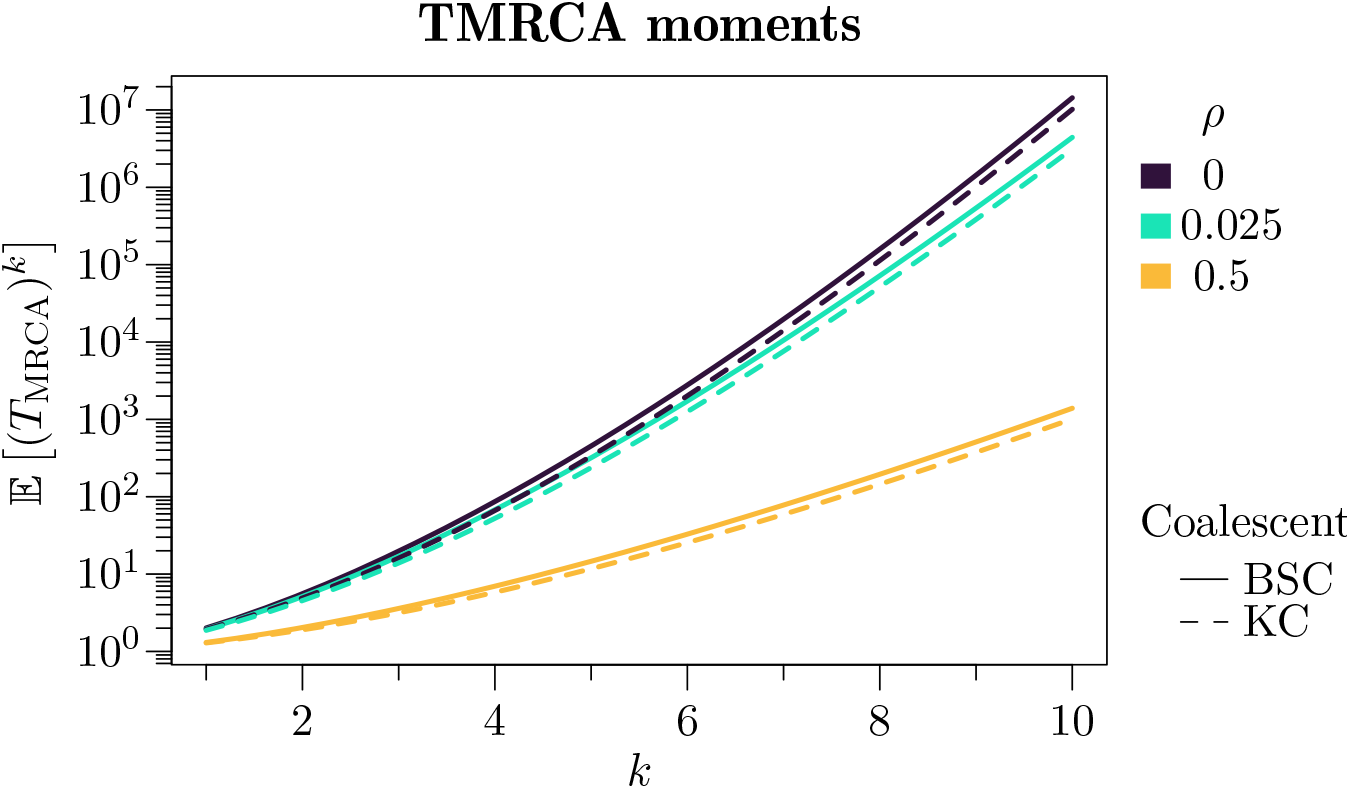
Moments of *T*_MRCA_ for different choices of order *k ∈* [1, 10] and exponential growth parameter *ρ*, for Kingman’s (KC) and Bolthausen-Sznitman coalescent (BSC) with *n* = 30.

Furthermore, in Figure 3 we demonstrate the use of our analytic formulas for the density of the random variable *T*_*MRCA*_ to compute the likelihood function, and its gradient, of a sample of *N* TMRCA’s. We use the latter to perform point estimation of *ρ ∈* [0, 1] and *α ∈* (0, 2], both jointly and separately, by finding the respective MLEs in each case. Even though obtaining closed expressions for the MLE is challenging, our explicit formulas for the likelihood function can be effectively used in parameter estimation through gradient-based optimization algorithms.

**Figure 3.**
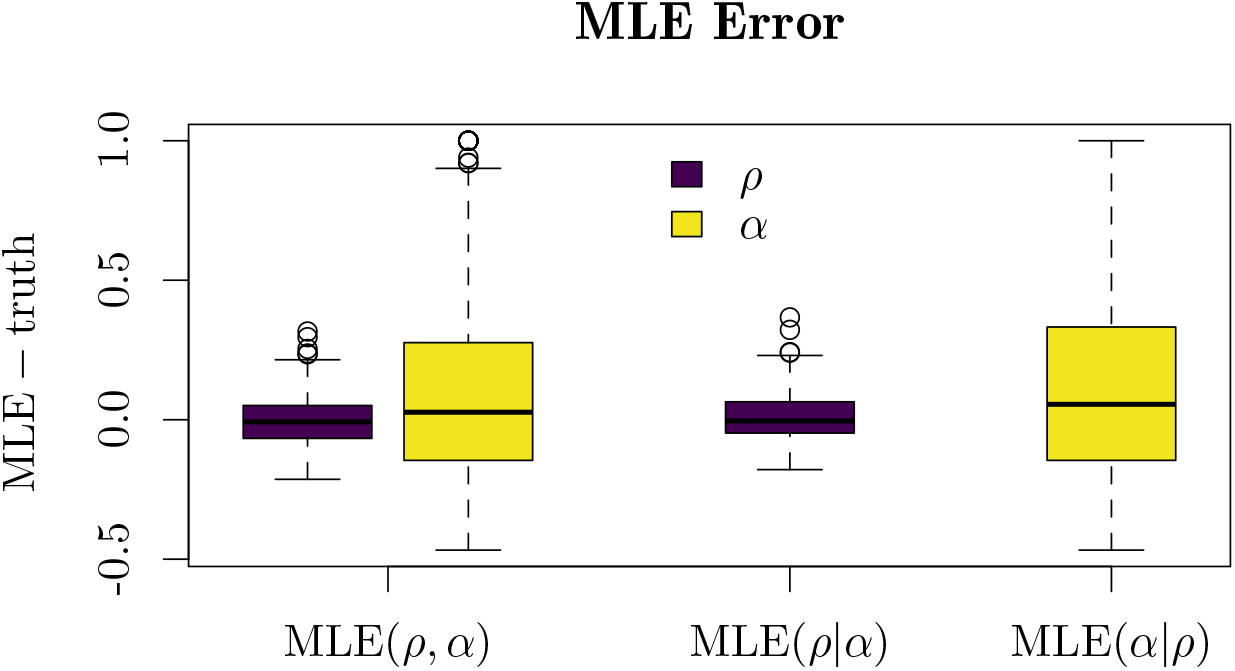
Boxplots showing the distribution of the difference between the estimated parameters (MLEs) and their true values *ρ* = 0.5 and *α* = 1 for 500 simulations of a sample of 50 TMRCAs of the Bolthausen-Sznitman coalescent with exponential growth, with sample size *n* = 30 individuals. On the left we show the differences of both parameters when both are simultaneously estimated, in the middle only the parameter *ρ* is estimated assuming that *α* = 1 is known, and on the right only the parameter *α* is estimated assuming *ρ* = 0.5 is known.

*Example 2: Populations with recurrent bottlenecks*. In Figure 4 we compare two models for populations undergoing recurrent bottlenecks, one homogeneous and one inhomogeneous. The first one consists of the symmetric coalescent introduced in [20]. The latter is a special case of Ξ-coalescents that arise from Wright-Fisher models that suffer from drastic decays of the population size for one generation; these decays occur at a constant rate. The symmetric coalescent is characterized by a function *F* on ℤ such that *F* (0) < ∞ and ∑ _*k ≥* 1_ *F* (*k*)/*k* < ∞; for convenience we also introduce a scalar parameter *A* > 0 modulating the overall coalescent rate. The dynamics are described as follows: when there are *b* blocks in the coalescent, at rate *AF* (*k*), we distribute the *b* blocks into *k* boxes uniformly at random, and blocks falling in the same box merge. This corresponds to setting 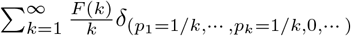. See also [43] for more general models of this type.

**Figure 4.**
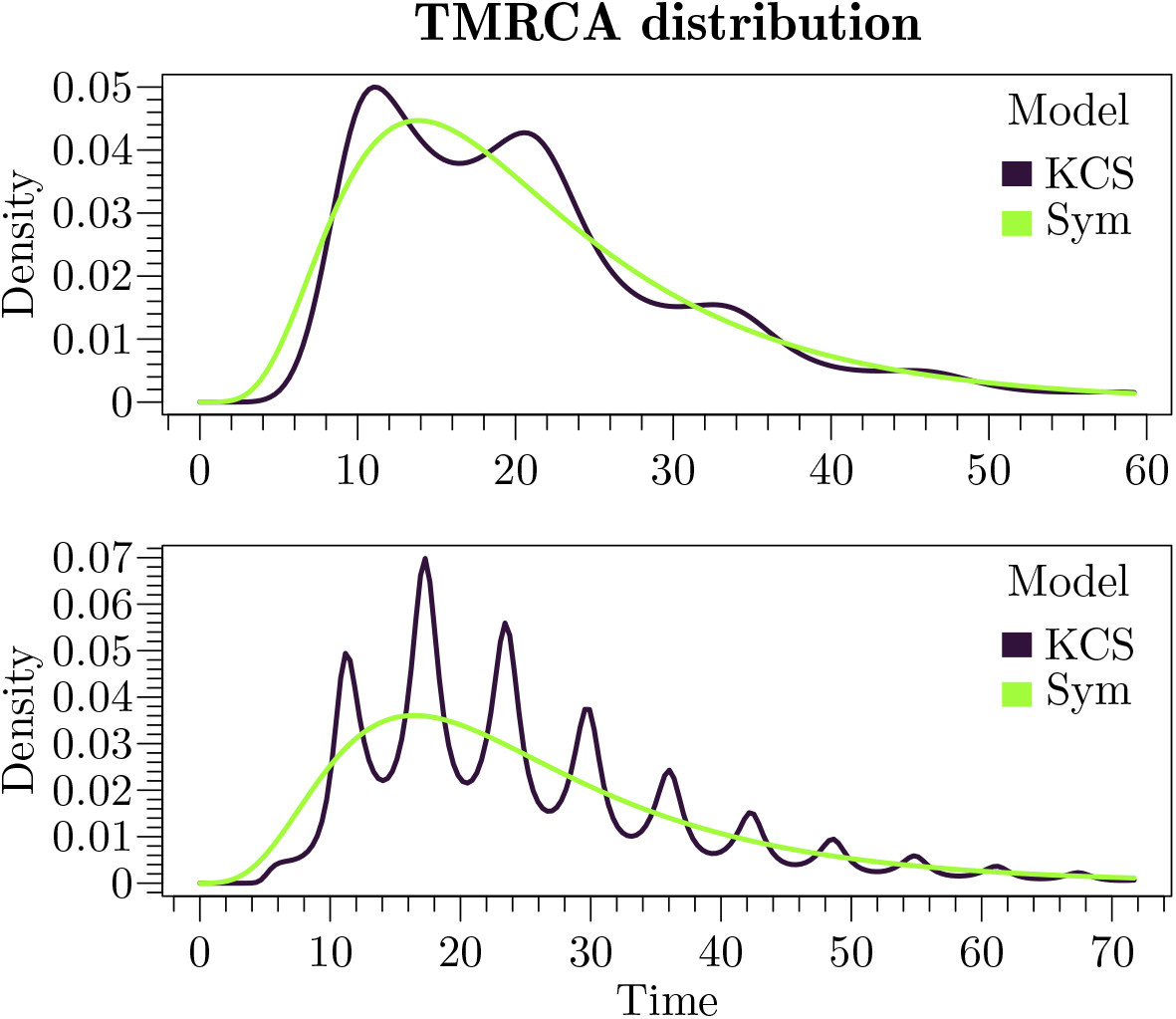
The density of *T*_MRCA_ for the two models with recurrent bottlenecks described in the main text, the sinusoidal Kingman’s coalescent (KCS) and the symmetric coalescent (Sym) with *n* = 30. Upper panel: *ε* = 0.8, *ω* = 0.5; lower panel: *ε* = 0.5, *ω* = 1. These parameters are chosen in order to show different behaviors of both the amplitude *ε* and the frequency 1*/ω* of the bottlenecks encoded by the function 1 + *ε* sin(*ωt*).

The second model for the genealogies of populations undergoing recurrent bottlenecks is the Kingman’s coalescent with sinusoidal time-change introduced in [14]. This model corresponds in our framework to setting *ζ*(*t*) = *B*(1 + *ε* sin(*ωt*)) where the parameter *B* > 0 gives the rate at which pair-wise merges occur, and *ε* and *ω* relate to the size and frequency of the bottleneck events.

In Figure 4 we compare the density of *T*_MRCA_ for the these two models. For the sinusoidal Kingman coalescent we fix two different combinations of *ε* and *ω*. For the symmetric coalescent we set *F* to be the density of a Poisson r.v. of parameter *λ* which we set to *λ* = *nε* where *n* is the initial number of blocks; whereas *A* is set to *A* = 1*/ω*. The choice of *λ* and *A*, respectively, heuristically match the amplitude and the frequency of the bottlenecks encoded by the term 1 + *ε* sin(*ωt*) in the sinusoidal Kingman’s coalescent. Finally, the remaining parameter *B* for the sinusoidal Kingman’s coalescent is chosen in order to ensure that the expectation 𝔼[*T*_MRCA_] is equal to that of the corresponding symmetric model (here, all the expectations 𝔼[*T*_MRCA_] are computed using (5) and they are matched numerically). The top of Figure 4 corresponds to the choice *ε* = 0.8 and *ω* = 0.5, whereas the bottom corresponds to *ε* = 0.5 and *ω* = 1. Note that, depending on the choice of the parameters *ε* and *ω*, the density of *T*_MRCA_ of the corresponding sinusoidal Kingman’s coalescent can be made to be multimodal, resembling the density of a discrete random variable. On the other hand, the density corresponding to the symmetric coalescent remains unimodal and appears as a continuous approximation of that of the sinusoidal Kingman’s case.

*Example 3: Populations with dormancy effect*. In this example we aim to study the effect of dormancy on the TMRCA of a sample of individuals. Two main genealogical models for populations with dormancy have been presented in the literature. The first model is a delayed coalescent [30, 22]. This model is obtained from Kingman’s coalescent by rescaling the length of all branches by a factor *β*^2^, where *β* is the expected number of generations one individual has to go backward in the past in order to find its parent, which is affected by dormancy. Thus, under this *weak seed bank* effect, *T*_MRCA_ can be obtained —in distribution— by simply multiplying *T*_MRCA_ of the classical Kingman’s coalescent by *β*^2^. Hence, its density *f*_*W*_ under the weak seed bank effect can be obtained from the density *f*_*K*_ of *T*_MRCA_ in Kingman’s coalescent via the formula

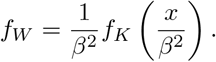

A second framework models populations where individuals can stay in a dormant state for longer periods of time [10, 21, 16]. This *strong seed bank* effect leads to a structured genealogical process, that differs from classical coalescents, where now lineages can be active or inactive. Active lineages can coalesce with regular Kingman dynamics, whereas inactive lineages do not coalesce. In addition, active lineages can turn into inactive lineages at rate 1 and vice versa. The relevant block-counting processes of the *strong seed bank* coalescent for a sample of size *n* is a two-dimensional process. It takes values of the form (*i, j*) with *i* + *j ≤ n*, where *i* denotes the number of active lineages and *j* the number of inactive lineages. Its starting configuration is (*n*, 0), which corresponds to index 1 in the matrix **S**, and its absorbing state is (1, 0). The process jumps from (*i, j*) to (*i* − 1, *j*) at rate 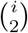, to (*i* − 1, *j* + 1) at rate *i* and to (*i* + 1, *j* − 1) at rate *j*. This Markov chain is compatible with the phase-type setting and the density of *T*_MRCA_ can thus be obtained as before by using the respective transition matrix **Q**(*t*), see [27].

In both cases, dormancy has a delaying effect on the TMRCA, increasing its value. We compare this effect with other delaying effects, such as exponential growth with a negative parameter, corresponding to genealogies of populations whose size decreases forward in time. As can be seen in (15), values of *ρ* < 0 can be readily used in the formula for the density. However the expectation is infinite in this case, because of the heavy tail of the distribution. All of the three effects (*weak seed bank, strong seed bank*, and exponential growth with negative parameter) are compared in Figure 5

**Figure 5.**
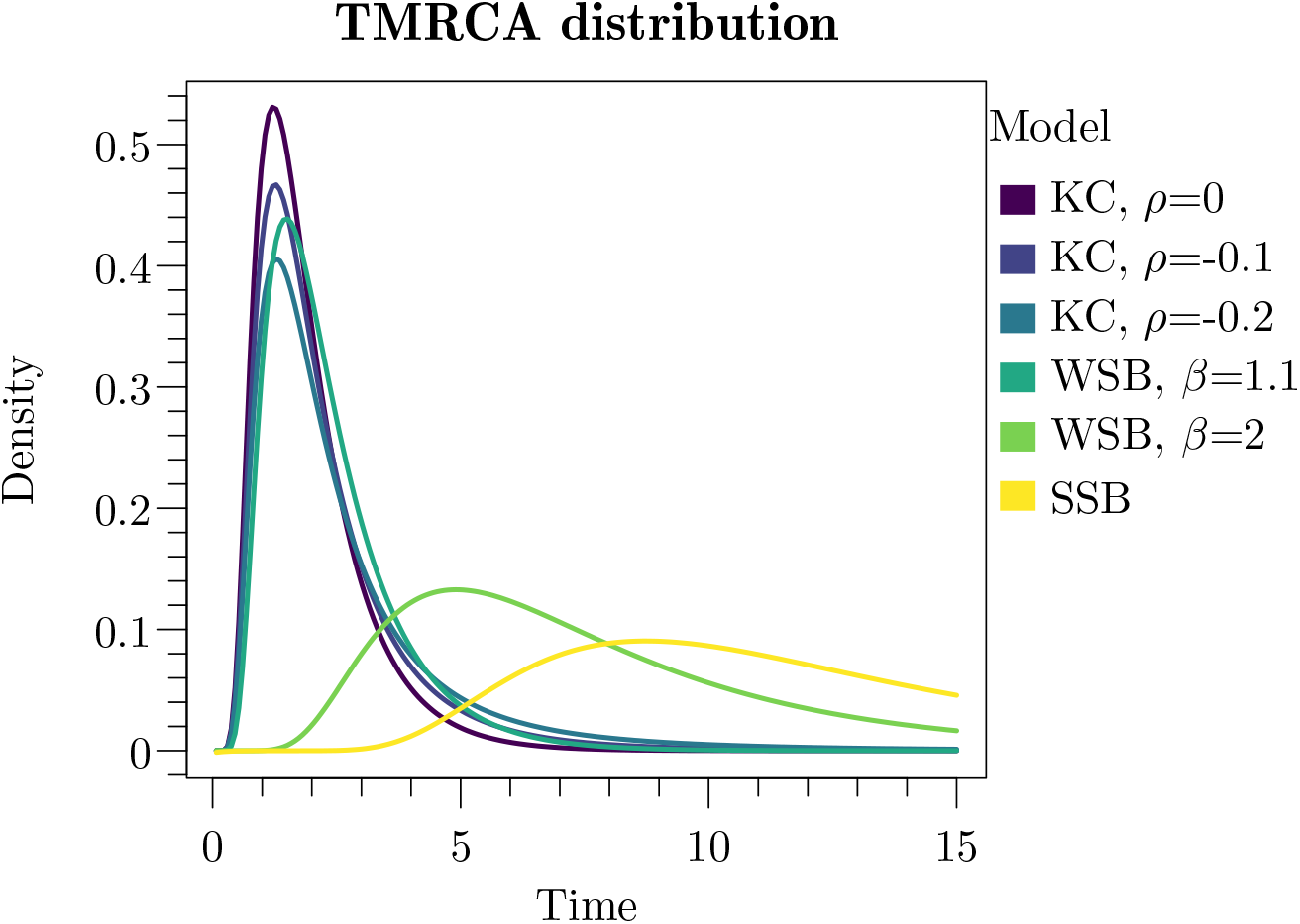
The density of *T*_MRCA_ with *n* = 30 for different delaying effects: exponential growth with negative parameters *ρ* = 0, −0.1, −0.2, *weak seed bank* effect (WSB) with *β* = 1.1, 2, and *strong seed bank* effect (SSB).

## 6 Discussion

In this manuscript, we exhibited a connection between general genealogical models of populations with variable size and the inhomogeneous phase-type theory described in [2]. We enriched this theory with interesting applications in population genetics, where the IPH theory provides explicit formulas for the TMRCA. In particular, we obtained expressions for its density and moments in a wide class of time-inhomogeneous coalescent processes. This improves previous results in the literature and also generalizes them to a much wider spectrum of models, including those that involve coalescents with simultaneous multiple mergers. This method is notably robust and can be applied to any Markovian genealogy starting with finitely many individuals. It also significantly eases the computational load present in inference applications by separating the effects of the time-inhomogeneity (the change in the population size captured by *ζ*(*t*)) and the coalescent dynamics.

This straightforward method does not readily generalize to other summary statistics in time-inhomogeneous coalescent models, such as the total branch length or the site frequency spectrum (SFS); or even the TMRCA in more complex time-inhomogeneous coalescent models, like the coalescent with recombination or the multispecies coalescent. On the one hand, it can be easily shown that these statistics are also IPH distributed; nonetheless, in these cases, the corresponding transition matrix is not easily factorized into a time-inhomogeneity and a coalescent component. For example, the density and moments of the total branch length can be expressed as a product integral of a time-dependent matrix for which computational methods must be developed. We note that the computation of this product integral can be recast in terms of PDE’s by adapting techniques of [37] or [7, Ch. 8.1.3]; however, the complexity and the substantially different nature of this approach make it fall out of the scope of this article.

Still, our method applies to a large family of genealogical models. For these models, the TMRCA has the double advantage of being the only statistic of a time-inhomogeneous coalescent process for which tractable formulas can be obtained, while also being able to distinguish between closely related population models. This makes the TMRCA a suitable tool for model selection and for the inference of the evolutionary dynamics of biological populations.

However, the study of the SFS for time-inhomogeneous coalescents motivates the development of a new theory for multivariate IPH distributions. This could also provide interesting insights into the covariance of the TMRCA and the total branch length. For now, this multivariate setting can only be established in a very restrictive form that does not cover many population genetics models (see [3]). These advances could also be essential in studying multivariate genealogical models such as recombination trees.

## Acknowledgement

This project was partially supported by DGAPA - PAPIIT grant IN102824, DGAPA - PASPA support. MS was supported by the National Institute of General Medical Sciences (NIGMS) of the National Institutes of Health under award R01GM146051. AHW was partially supported by the ANR project EPLER, and by the ANR LabEx CIMI (grant ANR-11-LABX-0040) within the French State Programme “Investissements d’Avenir.”

## References

[1] O.O. Aalen. Phase type distributions in survival analysis. Scandinavian Journal of Statistics, 22(4):447–463, 1995.

[2] H. Albrecher and M. Bladt. Inhomogeneous phase-type distributions and heavy tails. Journal of Applied Probability, 56(4):1044–1064, 2019.

[3] H. Albrecher, M. Bladt, and J. Yslas. Fitting inhomogeneous phase-type distributions to data: the univariate and the multivariate case. Scandinavian Journal of Statistics, 49(1):44–77, 2022.

[4] J. Bertoin. Random Fragmentation and Coagulation Processes. Number 102 in Cambridge Studies in Advanced Mathematics. Cambridge University Press, 2006.

[5] A. Bhaskar, Y.X. Wang, and Y. S. Song. Efficient inference of population size histories and locus-specific mutation rates from large-sample genomic variation data. Genome Research, 25(2):268–279, 2015.

[6] M. Birkner, H. Liu, and A. Sturm. Coalescent results for diploid exchangeable population models. Electronic Journal of Probability, 23:1–44, 2018.

[7] M. Bladt and B.F. Nielsen. Matrix-Exponential Distributions in Applied Probability. Springer, 2017.

[8] J. Blath, E. Buzzoni, J. Koskela, and M. Wilke Berenguer. Statistical tools for seed bank detection. Theoretical Population Biology, 132:1–15, 2020.

[9] J. Blath, M.C. Cronjäger, B. Eldon, and M. Hammer. The site-frequency spectrum associated with ξ-coalescents. Theoretical Population Biology, 110:36–50, 2016.

[10] J. Blath, A. González Casanova, N. Kurt, and D. Spano. The ancestral process of long-range seed bank models. Journal of Applied Probability, 50(3):741–759, 2013.

[11] E. Brunet and B. Derrida. Shift in the velocity of a front due to a cutoff. Physical Review E, 56(3):2597–2604, 1997.

[12] A. Cortines and B. Mallein. A n-branching random walk with random selection. Latin American Journal of Probability and Mathematical Statistics, 14(1):117–137, 2017.

[13] B. Eldon and J. Wakeley. Coalescent processes when the distribution of offspring number among individuals is highly skewed. Genetics, 172(4):2621–2633, 2006.

[14] A. Eriksson, B. Mehlig, M. Rafajlovic, and S. Sagitov. The total branch length of sample genealogies in populations of variable size. Genetics, 186(2):601–611, 2010.

[15] M. Fackrell. Modelling healthcare systems with phase-type distributions. Health Care Management Science, 12(1):11–26, 2008.

[16] M.C. Fittipaldi, A. González Casanova, and J.E. Nava. Lookdown construction for a moran seed-bank model. Electronic Communications in Probability, 29:1–14, 2024.

[17] F. Freund. Cannings models, population size changes and multiple-merger coalescents. Journal of Mathematical Biology, 80(5):1497–1521, 2020.

[18] Y. Fu and W. Li. Coalescing into the 21st century: An overview and prospects of coalescent theory. Theoretical Population Biology, 56(1):1–10, 1999.

[19] A. González Casanova, V. Miró Pina, E. Schertzer, and A. Siri-Jégousse. Asymptotics of the frequency spectrum for general dirichlet Ξ-coalescents. Electronic Journal of Probability, 29:1–35, 2024.

[20] A. González Casanova, V. Miró Pina, and A. Siri-Jégousse. The symmetric coalescent and Wright–Fisher models with bottlenecks. The Annals of Applied Probability, 32(1):235–268, 2022.

[21] A. González Casanova, L. Peñaloza, and A. Siri-Jégousse. The shape of a seed bank tree. Journal of Applied Probability, 59(3):631–651, 2022.

[22] A. González Casanova, L. Peñaloza, and A. Siri-Jégousse. Seed bank cannings graphs: How dormancy smoothes random genetic drift. Latin American Journal of Probability and Mathematical Statistics, 20(2):1165–1186, 2023.

[23] R.C. Griffiths and S. Tavaré. Sampling theory for neutral alleles in a varying environment. Philosophical Transactions: Biological Sciences, 344(1310):403–410, 1994.

[24] R.C. Griffiths and S. Tavaré. The age of a mutation in a general coalescent tree. Communications in Statistics. Stochastic Models, 14(1-2):273–295, 1998.

[25] J. Hein, M.H. Schierup, and C. Wiuf. Gene Genealogies, Variation and Evolution: A Primer in Coalescent Theory. Oxford University Press, 2005.

[26] A. Hobolth, I. Rivas-González, M. Bladt, and A. Futschik. Phase-type distributions in mathematical population genetics: An emerging framework. Theoretical Population Biology, 157:14–32, 2024.

[27] A. Hobolth, A. Siri-Jégousse, and M. Bladt. Phase-type distributions in population genetics. Theoretical Population Biology, 127:16–32, 2019.

[28] P. Hoscheit and O.G. Pybus. The multifurcating skyline plot. Virus Evolution, 5(2):vez031, 2019.

[29] I. Kaj and S.M. Krone. The coalescent process in a population with stochastically varying size. Journal of Applied Probability, 40(1):33–48, 2003.

[30] I. Kaj, S.M. Krone, and M. Lascoux. Coalescent theory for seed bank models. Journal of Applied Probability, 38(2):285–300, 2001.

[31] J. Kelleher, Y. Wong, A. W. Wohns, C. Fadil, P. K. Albers, and G. McVean. Inferring whole-genome histories in large population datasets. Nature Genetics, 51(9):1330–1338, 2019.

[32] J. F. C. Kingman. On the genealogy of large populations. Journal of Applied Probability, 19(A):27–43, 1982.

[33] S.M. Krone and C. Neuhauser. Ancestral processes with selection. Theoretical Population Biology, 51(3):210–237, 1997.

[34] S. Kumagai and M.K. Uyenoyama. Genealogical histories in structured populations. Theoretical population biology, 102:3–15, 2015.

[35] H. Li and R. Durbin. Inference of human population history from individual whole-genome sequences. Nature, 475:493–496, 2011.

[36] A.H. Marshall and S.I. McClean. Using Coxian phase-type distributions to identify patient characteristics for duration of stay in hospital. Health Care Management Science, 7(4):285–289, 2004.

[37] A. Miroshnikov and M. Steinrücken. Computing the joint distribution of the total tree length across loci in populations with variable size. Theoretical Population Biology, 118:1–19, 2017.

[38] M. Möhle. The coalescent in population models with time-inhomogeneous environment. Stochastic processes and their applications, 97(2):199–227, 2002.

[39] M. Möhle and S. Sagitov. Coalescent patterns in diploid exchangeable population models. Journal of Mathematical Biology, 47(4):337–352, 2003.

[40] R.A. Neher and O. Hallatschek. Genealogies of rapidly adapting populations. Proceedings of the National Academy of Sciences, 110(2):437–442, 2013.

[41] E. Schertzer and A.H. Wences. Genealogical transition in the noisy N-branching random walk. how stronger selection may promote genetic diversity. Preprint on arXiv: 10.48550/arXiv.2301.07762, 2023.

[42] J. Schweinsberg. Coalescent processes obtained from supercritical Galton–Watson processes. Stochastic Processes and their Applications, 106(1):107–139, 2003.

[43] A. Siri-Jégousse and A. H. Wences. Exchangeable coalescents beyond the Cannings class. Journal of Mathematical Biology, 90(1):1–40, 2024.

[44] L. Speidel, M. Forest, S. Shi, and S.R. Myers. A method for genome-wide genealogy estimation for thousands of samples. Nature Genetics, 51(9):1321–1329, 2019.

[45] J.P. Spence, J.A. Kamm, and Y.S. Song. The Site Frequency Spectrum for General Coalescents. Genetics, 202(4):1549–1561, 2016.

[46] F. Tajima. Evolutionary relationship of dna sequences in finite populations. Genetics, 105(2):437–460, 1983.

[47] G. Upadhya and M. Steinrücken. Robust inference of population size histories from genomic sequencing data. PLoS Computational Biology, 18(9):e1010419, 2022.

[48] A.W. Wohns, Y. Wong, B. Jeffery, A. Akbari, S. Mallick, R. Pinhasi, N. Patterson, D. Reich, J. Kelleher, and G. McVean. A unified genealogy of modern and ancient genomes. Science, 375(6583):eabi8264, 2022.

[49] K. Zeng, B. Charlesworth, and A. Hobolth. Studying models of balancing selection using phase-type theory. Genetics, 218(2):1–17, 2021.

[50] D. Živković and T. Wiehe. Second-Order Moments of Segregating Sites Under Variable Population Size. Genetics, 180(1):341–357, 2008.

